# The Shank/ProSAP N-terminal (SPN) domain of Shank3 regulates targeting to postsynaptic sites and postsynaptic signalling

**DOI:** 10.1101/2023.04.28.538665

**Authors:** Daniel Woike, Debora Tibbe, Fatemeh Hassani Nia, Victoria Martens, Emily Wang, Igor Barsukov, Hans-Jürgen Kreienkamp

## Abstract

Members of the Shank family of postsynaptic scaffold proteins (Shank1-3) link neurotransmitter receptors to the actin cytoskeleton in dendritic spines through establishing numerous interactions within the postsynaptic density (PSD) of excitatory synapses. Large Shank isoforms carry at their N-termini a highly conserved domain termed the Shank/ProSAP N-terminal (SPN) domain, followed by a set of ankyrin repeats. Both domains are involved in an intramolecular interaction which is believed to regulate accessibility for additional interaction partners, such as Ras family G-proteins, αCaMKII and cytoskeletal proteins. Here we analyse the functional relevance of the SPN-Ank module; we show that binding of active Ras or Rap1a to the SPN domain can differentially regulate the localization of Shank3 in dendrites. In Shank1 and Shank3, the linker between the SPN and Ank domains binds to inactive αCaMKII. Due to this interaction, both Shank1 and Shank3 exert a negative effect on αCaMKII activity at postsynaptic sites in mice *in vivo*. The relevance of the SPN-Ank intramolecular interaction was further analysed in primary cultured neurons; here we observed that in the context of full-length Shank3, a closed conformation of the SPN-Ank tandem is necessary for proper clustering of Shank3 on the head of dendritic spines. Shank3 variants carrying Ank repeats which are not associated with the SPN domain lead to the atypical formation of postsynaptic clusters on dendritic shafts, at the expense of dendritic spines. Our data show that the SPN-Ank tandem motif contributes to the regulation of postsynaptic signalling, and is also necessary for proper targeting of Shank3 to postsynaptic sites. Our data also suggest how missense variants found in autistic patients which alter SPN and Ank domains affect the synaptic function of Shank3.

## Introduction

Mutations in the *SHANK3* gene are associated with autism spectrum disorders (ASD) in human patients. Deletions, nonsense and splice site mutations have been reported, leading to a loss-of-function of one copy of *SHANK3*. In addition, a number of missense mutations have been found in patients with autism (1, 2). The encoded Shank3 protein forms, together with Shank1 and Shank2, a major class of scaffold proteins in the postsynaptic density (PSD) of excitatory, glutamatergic synapses. Shank proteins share a similar set of domains; this includes a ubiquitin-like domain at the N-terminus, which we have termed the Shank/ProSAP N-terminal (SPN) domain (3), followed by a set of ankyrin repeats (Ank), a Src homology 3 (SH3) domain, a PSD-95/DLG/ZO1 (PDZ) domain, a long proline-rich region and a C-terminal sterile alpha motif (SAM) domain (4). Via their PDZ domains, Shank proteins bind to members of the GKAP/SAPAP protein family. This allows for an indirect interaction between Shank proteins and the NMDA receptors through binding of SAPAPs to PSD-95, another postsynaptic scaffold protein (5, 6). On the other hand, several F-actin associated proteins bind to the proline-rich region (7-11), suggesting that Shank proteins connect different types of glutamate receptor complexes with signalling molecules and the actin cytoskeleton in the dendritic spine (12). The C-terminal SAM domain enables Shank2 and Shank3 (but not Shank1) to multimerize in a Zn^2+^-dependent manner. This is required for postsynaptic targeting of these two proteins and allows for cross-linking of multiple sets of Shank-associated protein complexes at postsynaptic sites (13-15).

In contrast to the rather well studied domains of Shank proteins mentioned above, the functional relevance of the N-terminal part (SPN and Ank domains) has remained unclear. By solving the three-dimensional structure of the N-terminus of Shank3 (residues 1-348, including SPN and Ank domains), we showed that the SPN domain adopts an ubiquitin-like (Ubl) fold which is similar to Ras binding domains (16). Subsequent studies confirmed that the SPN domain of Shank3 constitutes a high affinity binding site for several Ras family G-proteins, including H-Ras, KRas, Rap1a and Rap1b, and that there are actually two binding sites for Rap1 in the Shank3 N-terminus (16, 17). For the Ank repeats, Sharpin, α-fodrin, δ-catenin and HCN1 (Hyperpolarization Activated Cyclic Nucleotide Gated Potassium Channel 1) have been reported as interaction partners (18-21). In addition, the SPN domain interacts with the Ank repeats in an intramolecular manner and limits access to the Ank repeats for some of its interaction partners (3, 16).

Several recent studies have begun to shed light on the relevance of the intramolecular interaction between SPN and Ank. Thus, it was observed that the linker region between both domains, and part of the SPN domain constitute a binding interface for the α-subunit of the calcium-/calmodulin dependent kinase II (αCaMKII). The αCaMKII protein must be in its non-phosphorylated, inactive form, and the SPN-Ank tandem must be in a closed conformation for binding to occur (22, 23). On the other hand, F-actin binds to the SPN domain when the SPN-Ank tandem is in an open conformation (24). It is currently not clear whether some kind of regulated opening of the closed SPN-Ank conformation occurs *in vivo*. Interestingly, several patient derived missense mutations lead to such an opening, either through unfolding of the SPN domain (by the L68P variant) or by disrupting the contact area on the side of the Ank domain (by the P141A variant) (3, 23, 25).

Here we analysed in detail the relevance of the SPN domain for Shank3 localization and function. We observed that binding of small G-proteins may direct the localization of Shank3 in dendrites. We show that small G-proteins and the αCaMKII may bind simultaneously to the folded Shank3 N-terminus. Due to this interaction, Shank3 regulates the activity of αCaMKII at postsynaptic sites. Finally, we show that the SPN-Ank interaction is required to prevent formation of irregular Shank3 clusters on the dendritic shaft.

## Materials and Methods

### Expression constructs

cDNA coding for SPN+Ank domains of human Shank1 (residues 72-555; NM_016148.5) and human Shank2 (residues 1-423; database NM_012309.5) were PCR amplified using primers carrying appropriate restrictions sites, and cloned into pmRFP-N vectors (Clontech). A construct coding for N-terminally GFP-tagged full-length rat Shank3 in the pHAGE vector was obtained from Alex Shcheglovitov (University of Utah), and has been used before (16). Deletion constructs (ΔSPN; ΔSPN+Ank) were generated by introducing SalI restriction sites at appropriate positions using site directed mutagenesis, followed by cutting out a SalI/SalI fragment and religating the remaining vector. Rat Shank3 deletion constructs in pmRFP-N3 (Clontech) were described before (23). Numbering of Shank3 residues is based on Uniprot entry Q9JLU4. Mutations were introduced using Quik-Change II site-directed mutagenesis kit (Agilent), using two complementary oligonucleotides carrying the mutated sequence. Constructs coding for HA-tagged HRas G12V and Rap1a G12V were obtained from Georg Rosenberger (UKE Hamburg, Germany). A construct for expression of N-terminally T7-tagged αCaMKII was described before (23).

### Cell culture and transient transfection

Human embryonic kidney (HEK) 293T cells were cultured in Dulbecco’s Modified Eagle Medium supplemented with 10% fetal bovine serum and 1% Penicillin/Streptomycin. Transient transfection of 293T cells was performed using Turbofect Transfection Reagent (Thermo Scientific) according to the manufacturer’s instructions.

### Cell lysis and immunoprecipitation

Cell lysis was performed using immunoprecipitation (IP) buffer (50 mM Tris-HCl, pH 8, 120 mM NaCl, 0.5% NP40, 1 mM EDTA). Lysates were centrifuged at 20,000 x g for 15 min at 4 °C. Immunoprecipitation from supernatants was then performed using 20 μl of RFP-trap beads (Chromotek, Munich, Germany; 2 hrs at 4 °C on a rotator). Precipitates were washed in IP buffer and then processed together with input samples for SDS-PAGE and Western blotting.

### SDS-PAGE and Western blot

Proteins were separated on SDS-PAGE under denaturing conditions and transferred to nitrocellulose membrane using a MINI PROTEAN II^™^ system (Bio-Rad). Membranes were blocked with 5% milk powder/TBS-T and incubated with the primary antibodies overnight at 4 °C, followed by washing in TBS-T and then HRP-linked secondary antibodies at room temperature for 1 h. Membranes were scanned using a ChemiDoc^™^ MP Imaging System (Bio-Rad) and images were processed and further analysed using Image Lab Software (Bio-Rad).

### Animals

For preparing primary neuronal cultures, brain tissue was isolated from Rattus norvegicus embryos. Pregnant rats (Envigo; 4-5 months old) were sacrificed on day E18 of pregnancy using CO2 anaesthesia, followed by decapitation. Neurons were prepared from all embryos present, regardless of gender (12-16 embryos). All animal experiments were performed in compliance with ARRIVE guidelines. Shank1 and Shank3 ko mouse lines were obtained from Carlo Sala (CNR; Milano, Italy) and Tobias Böckers (Univ. of Ulm, Ulm, Germany), respectively. All animal experiments were approved by the Behörde für Justiz und Verbraucherschutz, Freie und Hansestadt Hamburg and conducted in accordance with the guidelines of the Animal Welfare Committee of the University Medical Center (Hamburg, Germany) under permission numbers Org766, Org1088 (rats) and N19/2022 (mice).

### Preparation of postsynaptic density from mouse hippocampus and cortex

For preparation of PSDs, animals were sacrificed using CO_2_ anaesthesia, followed by decapitation. Hippocampi and cortices were dissected from brains. Tissue from five animals was pooled for a single preparation. PSD fractions were prepared by series of centrifugation and ultracentrifugation steps, as previously described (21, 26, 27).

### Neuron culture and transfection

For preparing primary neuron cultures, hippocampal tissue was isolated from *Rattus norvegicus* embryos. Pregnant rats (Envigo; 4-5 months old) were sacrificed on day E18 of pregnancy using CO_2_ anaesthesia, followed by decapitation. Neuron cultures were prepared from all embryos present, regardless of gender (14-16 embryos). The hippocampal tissue was dissected, and hippocampal neurons were extracted by trypsin digestion, followed by mechanical dissociation. Cells were plated on glass cover slips and cultivated in Neurobasal medium supplemented with 2% B27, 1% Glutamax and 1% Penicillin/Streptomycin. Neurons were transfected after seven days *in vitro* (DIV7) using the calcium phosphate method. All animal experiments were approved by, and conducted in accordance with, the guidelines of the Animal Welfare Committee of the University Medical Center (Hamburg, Germany) under permission numbers Org766 and Org1018.

### Immunocytochemistry (ICC)

Neurons were fixed (DIV14), with 4% paraformaldehyde in PBS and permeabilized with 0.1% Triton X-100 in PBS for 5 min at room temperature. After blocking (10% horse serum in PBS) for 1 h at room temperature, cells were incubated with corresponding antibody overnight followed by washing and then 1 h of incubation with Alexa Fluor coupled secondary antibodies. The coverslips were mounted onto glass microscopic slides using ProLong^™^ Diamond Antifade mounting medium.

### Microscopy

Confocal images were acquired with a Leica Sp5 confocal microscope using a 63x objective. Quantitative analysis for images was performed using ImageJ. Primary dendrites were counted at a ring within 10 µm distance from the neuronal cell body. The counting of clusters along dendritic branches was performed using the Multi-Point tool of ImageJ.

### Antibodies

The following primary antibodies were used: mouse anti-GFP (Covance MMS-118P-500, RRID:AB_291290; WB: 1:3000); rat anti-RFP (Chromotek 5F8ChromoTek 5f8-100, WB: 1:1000); chicken anti-MAP2 (antibodies Antibodies-Online ABIN361345, ICC: 1:1000); mouse anti PSD-95 (Thermo Fisher MA1-046; ICC: 1:500); rabbit anti-HA (Sigma Aldrich #H9658 ICC 1:200); rabbit anti-CaMKIIα (abcam ab52476) and rabbit anti-CaMKIIα phospho-T286 (abcam ab32678). HRP-labeled goat secondary antibodies were purchased from Jackson ImmunoResearch and used for WB at 1:2500 dilution. For ICC, Alexa 633 goat anti-chk IgG (Invitrogen A21103), Alexa 633 goat anti-mouse IgG (Invitrogen A21050), Cy3 goat anti-rabbit IgG (abcam ab6939) and Alexa 405 goat anti-chk IgG (abcam ab175675) were used at 1:1000 dilution.

### DSF measurements

Prepapration of His_6_-SUMO tagged Shank3 fusion proteins and measurement of differential scanning fluorimetry DSF was performed as described (23).

### Evaluation of data

Statistical significance was determined using Prism8 software (GraphPad, San Diego, CA) and analysed by Student’s t-test or one-way ANOVA with post hoc Dunnett’s or Tukey’s test. All data are presented as mean ± SD.

## Results

### Active Ras family G-proteins control localization of Shank3 in hippocampal neurons

In neurons, Shank proteins are targeted to the postsynaptic core complex through direct interaction with SAPAP proteins which links Shank3 to PSD-95 and NMDA receptors, and through multimerization of the C-terminal SAM domains (28, 29). We analyzed the effect of active G-proteins on the localization of Shank3 by coexpression of Shank3 with active forms of Rap1 or HRas (Rap1a-G12V; HRas-G12V) in primary hippocampal neurons. Using confocal imaging, we observed that GFP-tagged Shank3 localizes to dendritic protrusions in a clustered manner, in keeping with the known targeting of Shank3 to dendritic spines (Fig. 1, upper panel). When coexpressed with Rap1-G12V, this localization did not change; by staining for the HA-tagged Rap1, we determined that Rap1 colocalizes with GFP-Shank3 in dendritic spines (Fig. 1, lower panel). Upon coexpression with HRas-G12V, this pattern changed significantly (Fig.1, middle panel). The active HRas-G12V protein shows a membrane-associated localization, in agreement with the known targeting of HRas to membranes via the palmitoyl- and farnesyl lipid anchors (30, 31). Along dendrites, HRas appeared to be localized along dendritic shafts, but did not noticeably extend into dendritic protrusions or spine–like structures. Importantly, HRas-G12V was found to be highly colocalized with GFP-Shank3 along dendritic shafts.

**Figure 1.**
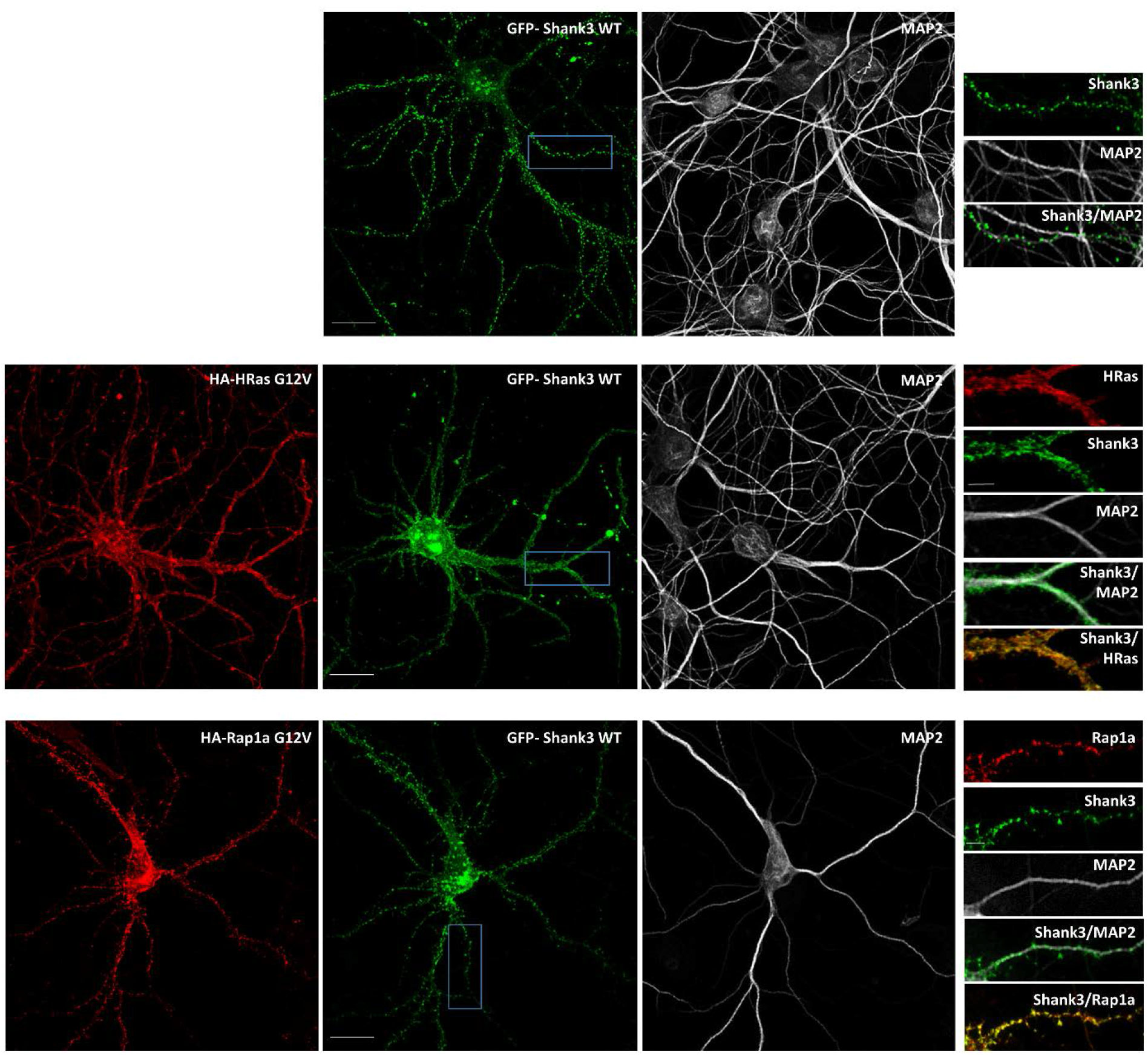
Active HRas affects the localization of Shank3 in primary cultured neurons. Primary hippocampal neurons were transfected with GFP-Shank3 alone (upper panels) or together with HA-tagged active (G12V mutant) forms of either HRas or Rap1a. Neurons were stained for HA (red) and the dendritic marker MAP2 (gray). Left panels show overview images of neurons (scale bar 20 μm) and right panels show the magnified boxed area (scale bar 5 μm). The overexpressed constitutively active form of HRas (G12V) shows a membrane-associated pattern along dendritic shafts, whereas active form of Rap1a shows a higher signal intensity in dendritic spines. In both cases Shank3 WT is highly colocalized with the active, GTP-bound form of the small G-proteins.

Further investigations using the SPN mutant variants L68P and R12C showed that these two mutants, which are deficient in HRas binding, did not noticeably change their localization upon coexpression with HRas-G12V (Fig. 2A). Thus, both variants remained clustered in dendritic spines, and we observed reduced colocalization with the active HRas-G12V when compared to WT Shank3 (Fig. 2B). These data indicate that binding to active HRas via its SPN domain alters the localization of WT Shank3, but not of mutant forms of Shank3 which cannot bind to active HRas.

**Figure 2.**
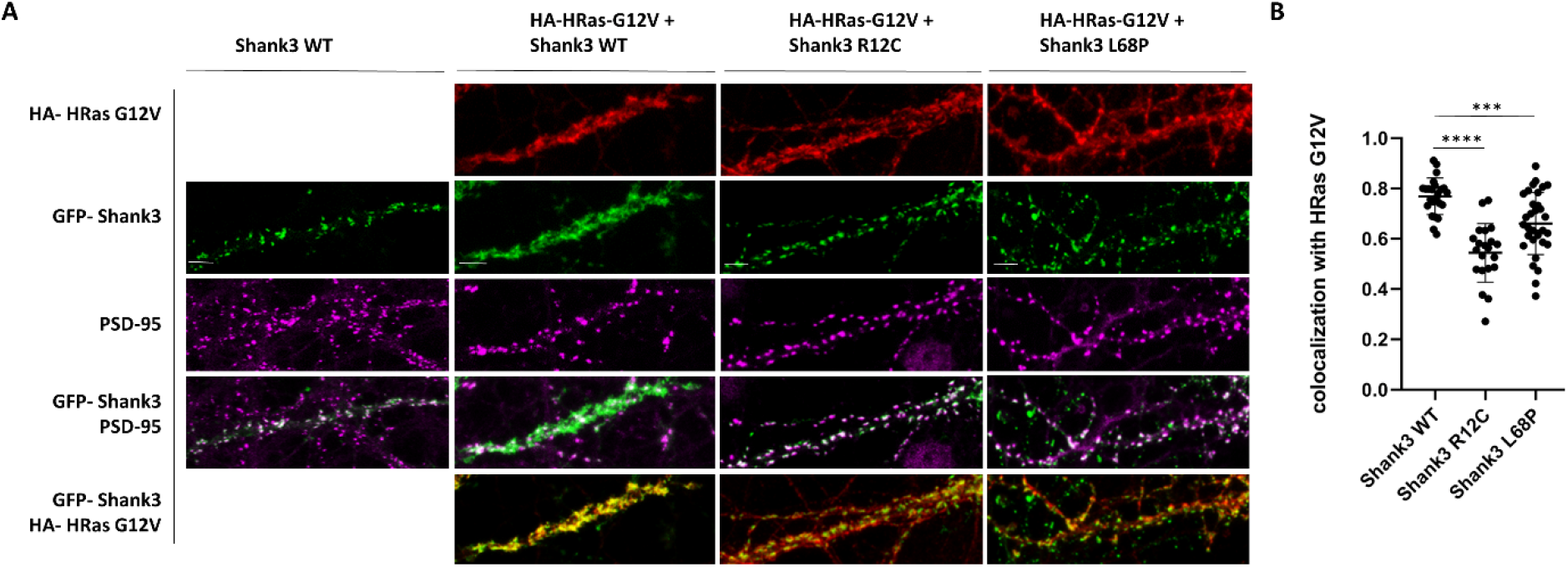
Localization pattern of Shank3 variants in the presence of HRas in cultured neurons. **A**. Neurons overexpressing active HA-HRas together with GFP-tagged Shank3, either WT or SPN mutants L68P and R12C, were stained for HA (red) and endogenous postsynaptic marker PSD-95 (magenta). In the presence of active HRas, the overexpressed Shank3 WT shows a high degree of colocalization with active G12V HRas and subsequently changes in the synaptic clustering pattern whereas mutant variants of Shank3 (R12C and L68P) show a punctate postsynaptic pattern highly colocalized with endogenous PSD-95 (scale bar 5 μm). **B**. To analyse the colocalization of Shank3 variants with active HRas, the Pearson correlation coefficient (PCC) of green and red channels of various dendritic pieces from 10-12 neurons (obtained from 4 independent experiments) was measured using the JACoP plugin of the ImageJ software. Data show a significantly higher degree of colocalization between active HRas and Shank3 WT compared to the mutant variants of Shank3. ***,****: significantly different, p<0.001, 0.0001, respectively. Data from four independent experiments; one-way ANOVA followed by Dunnett’s multiple comparisons test.

The altered localization of Shank3 upon expression of active HRas could be due to the loss of postsynaptic specializations in spines. To determine whether this is the case, transfected neurons were stained for the endogenous postsynaptic marker PSD-95. Here we observed that PSD-95 is still present in a clustered manner at the dendritic sites of HRas-G12V expressing neurons with a typical distribution pattern (Fig. 2A). SPN domain mutants, but not WT Shank3, were highly colocalized with PSD-95 in the presence of an overexpressed active form of HRas. Thus, the presence of active HRas does not interfere with the formation of dendritic spines and the postsynaptic density but leads to a selective absence of WT GFP-Shank3 from postsynaptic sites.

### Small G-proteins and the αCaMKII do not compete for binding to the Shank3 N-terminus

The N-terminus of Shank3 (SPN+Ank domains) binds not only to active Ras variants, but also to the inactive form of αCaMKII (22, 23). By coexpression/coimmunprecipitation of mRFP-tagged Shank fragments with T7-tagged αCaMKII, we observed here that Shank3, and to a lesser extent Shank1 bind to αCaMKII, whereas almost no binding could be detected for Shank2 (Fig. 3).

**Figure 3.**
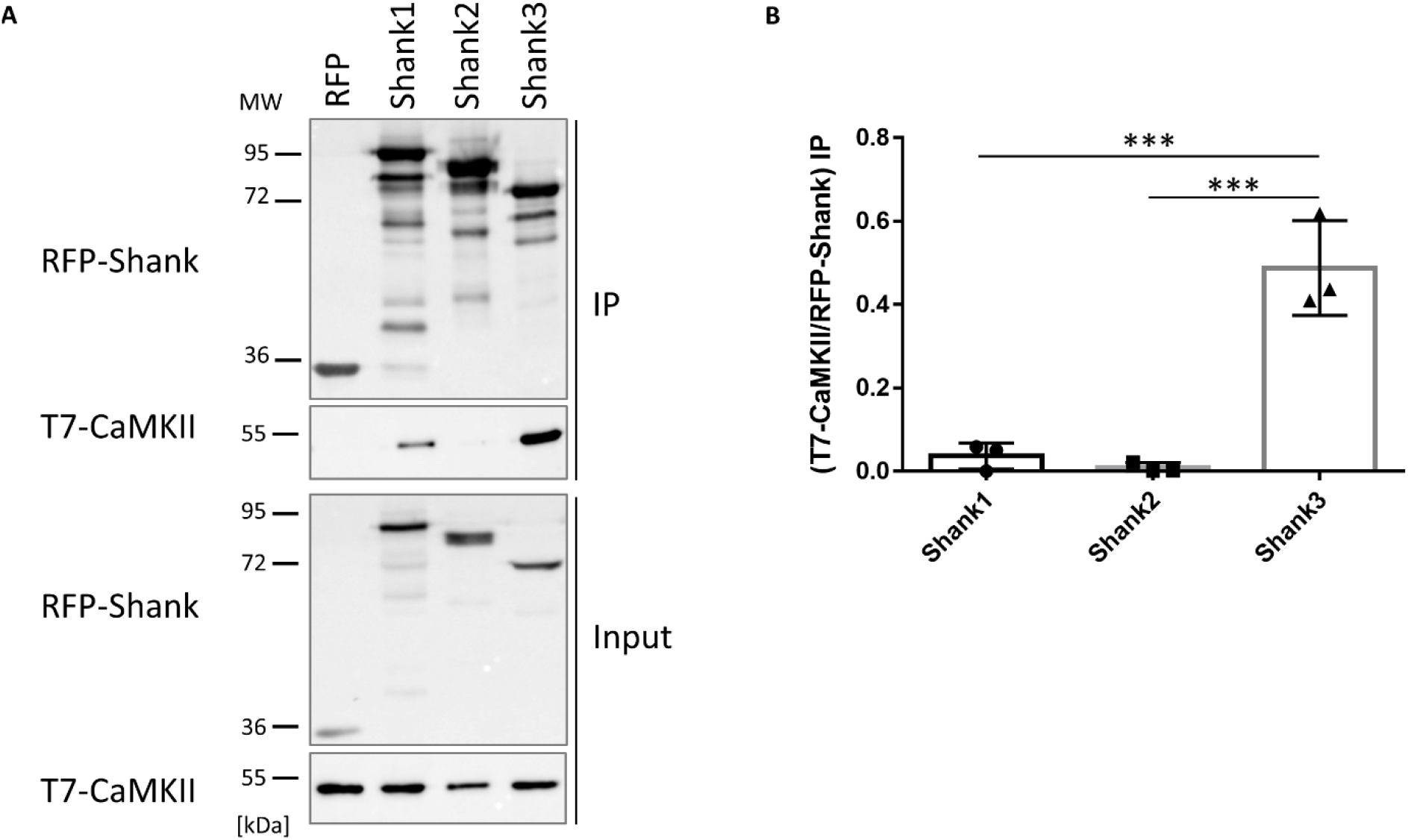
Shank isoforms differentially interact with αCaMKII. **A**. RFP-tagged N-termini (SPN+Ank domains) of different Shank isoforms, or mRFP alone, were coexpressed in 293T cells with T7-tagged αCaMKII. After cell lysis, RFP-tagged proteins were immunoprecipitated using the mRFP-trap matrix. Input and precipitate (IP) samples were analysed by Western blotting using mRFP- and T7-specific antibodies. **B**. Quantitative analysis. Signal intensities in IP samples for T7-αCaMKII were divided by IP signals for mRFP-Shank isoforms. ***, significantly different, p<0.001, respectively; data from three independent experiments; one-way ANOVA, followed by Tukey’s multiple comparisons test.

Interactions with Ras/Rap G-proteins and with αCaMKII are dependent on the presence of Arg12 in the Shank3 SPN domain (16, 22). To determine whether this leads to competition between these two signaling molecules, we expressed αCaMKII and active (G12V-mutant) forms of HRas or Rap1a with the mRFP-tagged Shank3 N-terminus. Upon immunoprecipitation of Shank3 using the mRFP-trap matrix, we observed that both the small G-proteins and αCaMKII were coprecipitated efficiently with Shank3. As expression of small G-proteins did not affect αCaMKII binding, and expression of αCaMKII did not affect Ras or Rap binding, we conclude that αCaMKII does not compete with these G-proteins for binding to Shank3 (Fig. 4).

**Figure 4.**
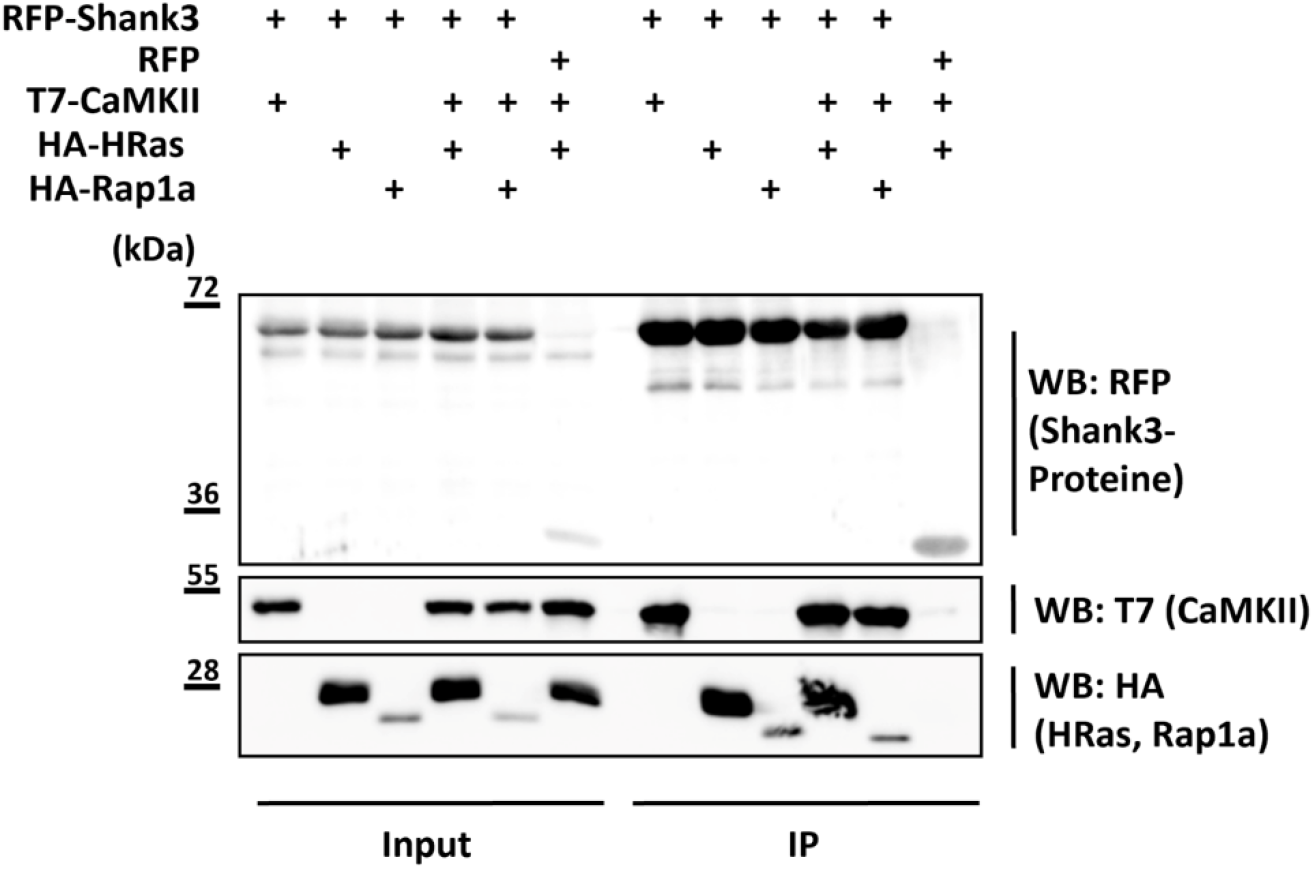
The αCaMKII does not compete with small G-proteins for binding to the Shank3 N-terminus. 293T cells expressing mRFP-tagged Shank3 N-terminus (SPN+Ank), or mRFP control, in various combinations with T7-tagged αCaMKII and active G-proteins were lysed, followed by immunoprecipitation of mRFP-tagged proteins. Input and IP samples were analysed by Western blotting using the antibodies indicated.

### Shank isoforms carrying the SPN domain control postsynaptic αCaMKII activity

Interactions with inactive αCaMKII and active Ras variants suggests that the N-terminal part of Shank3 may be involved in the regulation of postsynaptic signal transduction. In the HEK293T cell model, expression of Shank3, but not of its R12C and L68P variants, inhibits HRas-G12V mediated activation of the MAP kinase pathway (Fig. 5A, B). However, in lysates from cortex or hippocampus of Shank3 deficient mice, we did not observe a significant difference in activity of Erk_1/2_, or the Akt kinase, two major targets of Ras signaling in neurons (Fig. 5C-F).

**Figure 5.**
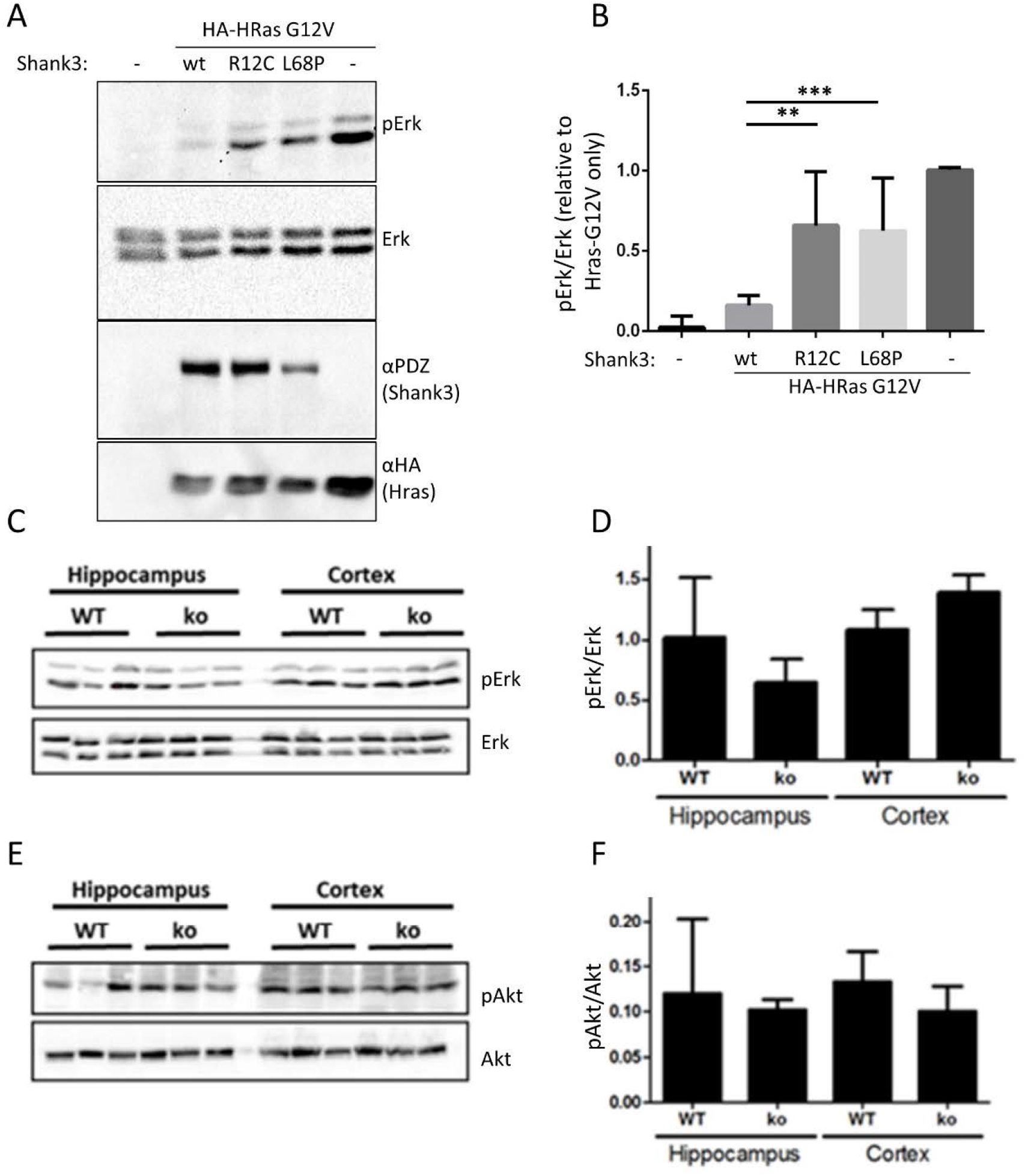
Role of the Shank3 SPN domain in the MAP kinase pathway. **A**. 293T cells were transfected with vectors coding for HA-tagged HRas-G12V and full length Shank3 (WT or mutants). Cell lysates were analysed by Western blotting using the antibodies indicated. **B**. Quantitative analysis of the data shown in A. The ratio of pErk to total Erk signal was calculated and normalized to the value obtained in the absence of Shank3 coexpression. **C**. Hippocampal and cortical lysates were prepared from WT and Shank3 ko mice. Samples containing equal amounts of protein were analysed by Western blotting using pErk and Erk specific antibodies. **D**. Quantitative analysis of the data shown in C; data are plotted as the ratio of pErk to Erk signal. **E**. Samples shown in C were analysed by Western blotting using pAkt and Akt specific antibodies. **F**. Quantitative analysis of the data shown in E; data are plotted as the ratio of pAkt to Akt signal. **, ***; significantly different from WT condition, p<0.01, 0.001, respectively; one-way ANOVA, followed by Dunnett’s multiple comparisons test; n=10. For D and F, no significant differences between WT and ko conditions were determined using a Student’s t-test (n=3).

For the αCaMKII, we also analysed the postsynaptic density fraction as αCaMKII is enriched at the synapse. We prepared the postsynaptic density fractions from mice lacking either Shank1 or Shank3. In the case of Shank1-ko, this is a complete knockout as described initially by (32). In the case of Shank3, isoforms containing the N-terminal SPN and Ank domains are missing due to the deletion of exon 11, but shorter isoforms initiating at the PDZ domain are present (33). PSD fractions were analysed by Western blotting using αCaMKII and phospho-T286 αCaMKII specific antibodies. Here we observed that in both mouse strains, the amount of αCaMKII in the PSD was not altered; however, the fraction of active (phosphorylated at T286) αCaMKII was significantly increased in hippocampus and cortex of Shank3 deficient mice, and in the cortex of Shank1 ko mice (Fig. 6).

**Figure 6.**
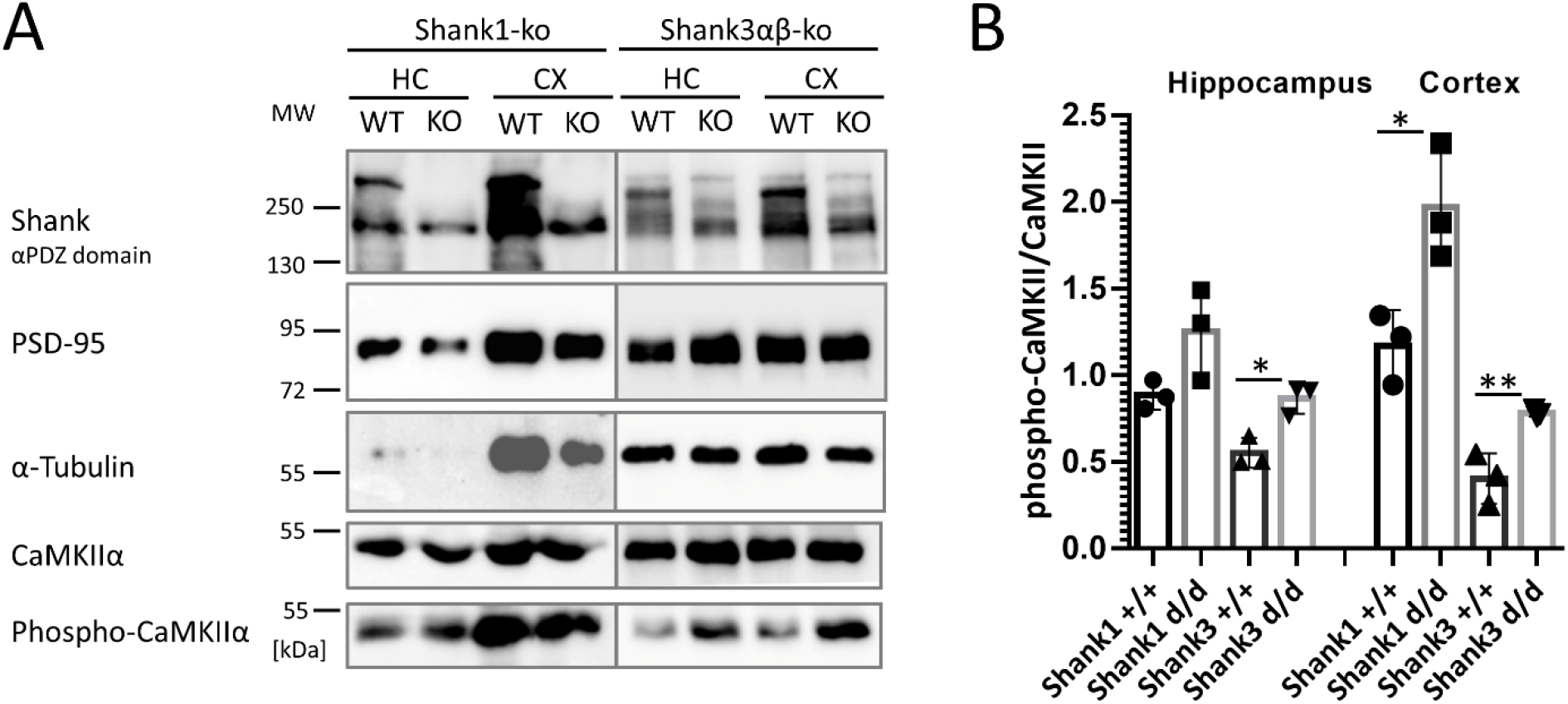
Shank proteins with N-terminal SPN and Ank domains regulate postsynaptic αCaMKII activity. **A**. The postsynaptic density fraction was prepared from hippocampus and cortex of Shank1 and Shank3 deficient mice. Samples were analyzed by Western blotting using the antibodies indicated. For Shank1, an anti-PDZ domain antibody was used which recognizes also Shank2, and to a lesser degree Shank3. For Shank3 samples, a Shank3-specific antibody was used. Note that shorter variants of Shank3 lacking SPN and Ank domains are still expressed, whereas large isoforms running at 250 kDa are lost. **B**. Quantification of the αCaMKII data shown in A. The ratio between phospho-CaMKII to pan-CaMKII was calculated. *,**: significantly different, p<0.05, 0.01, respectively. Data from three independent experiments; Student’s t-test.

### A closed conformation of the Shank3 N-terminus is required for the formation of spine associated synapses

For further functional analysis, we generated constructs of Shank3 containing the full–length coding sequence but lacking the SPN domain (ΔSPN), or lacking the SPN and Ank domains (ΔSPN+Ank) and transfected those into primary cultured hippocampal neurons. Neurons were fixed and stained for MAP2 and the postsynaptic marker PSD-95. As seen before, WT Shank3 was localized to dendritic spines where it perfectly colocalized with PSD-95 (Figs. 7, 8A). In contrast, the protein expressed from the ΔSPN deletion construct was in many cases found in proximity, or immediately adjacent to the main dendritic shaft. In addition, about 10% of dendritic Shank3 clusters were found on long, thin protrusions from the dendrite. All of these clusters were colocalized with PSD-95. In additional experiments, staining with the presynaptic marker vGlut1 showed that Shank3 clusters indeed represented synaptic contacts carrying a pre- and a postsynapse (Fig. 8B). With the ΔSPN+Ank construct, lacking the complete N-terminus, the localization of the expressed GFP-Shank3 protein returned to a “normal” pattern, as the number of clusters adjacent to the dendritic shaft was largely reduced, and most Shank3 clusters were localized on dendritic spines (Fig. 7,8). Colocalization with both PSD-95 and vGlut1 was observed, suggesting that regular spine-associated synapses have been formed (Fig. 8A, B).

**Figure 7.**
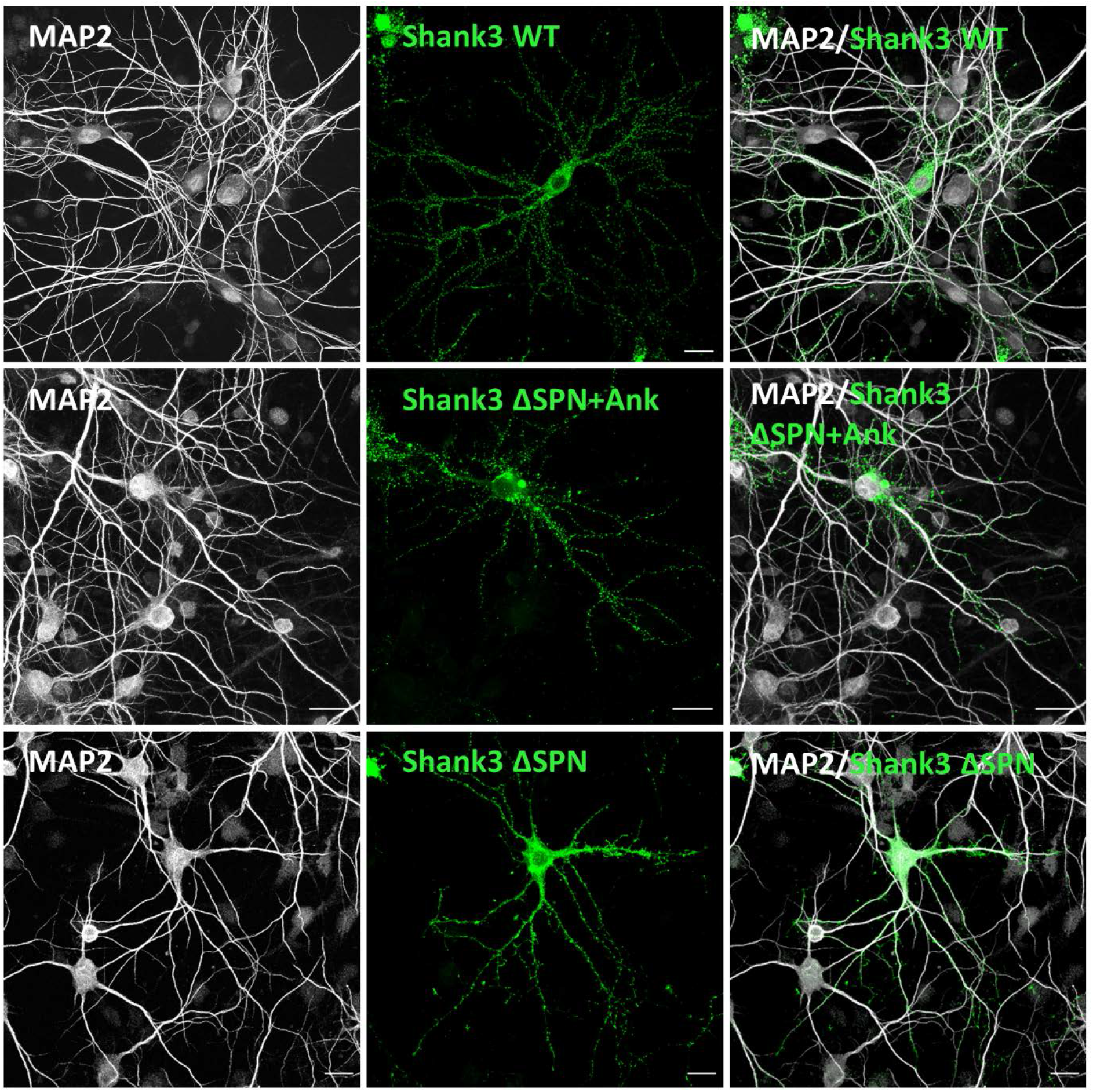
Expression of N-terminal Shank3 deletion constructs in neurons. Primary cultured hippocampal neurons were transfected with constructs coding for full-length WT Shank3 or the ΔSPN+Ank, or ΔSPN constructs, as indicated. Neurons were fixed and costained for the dendritic marker MAP2 (scale bar 20 µm).

**Figure 8.**
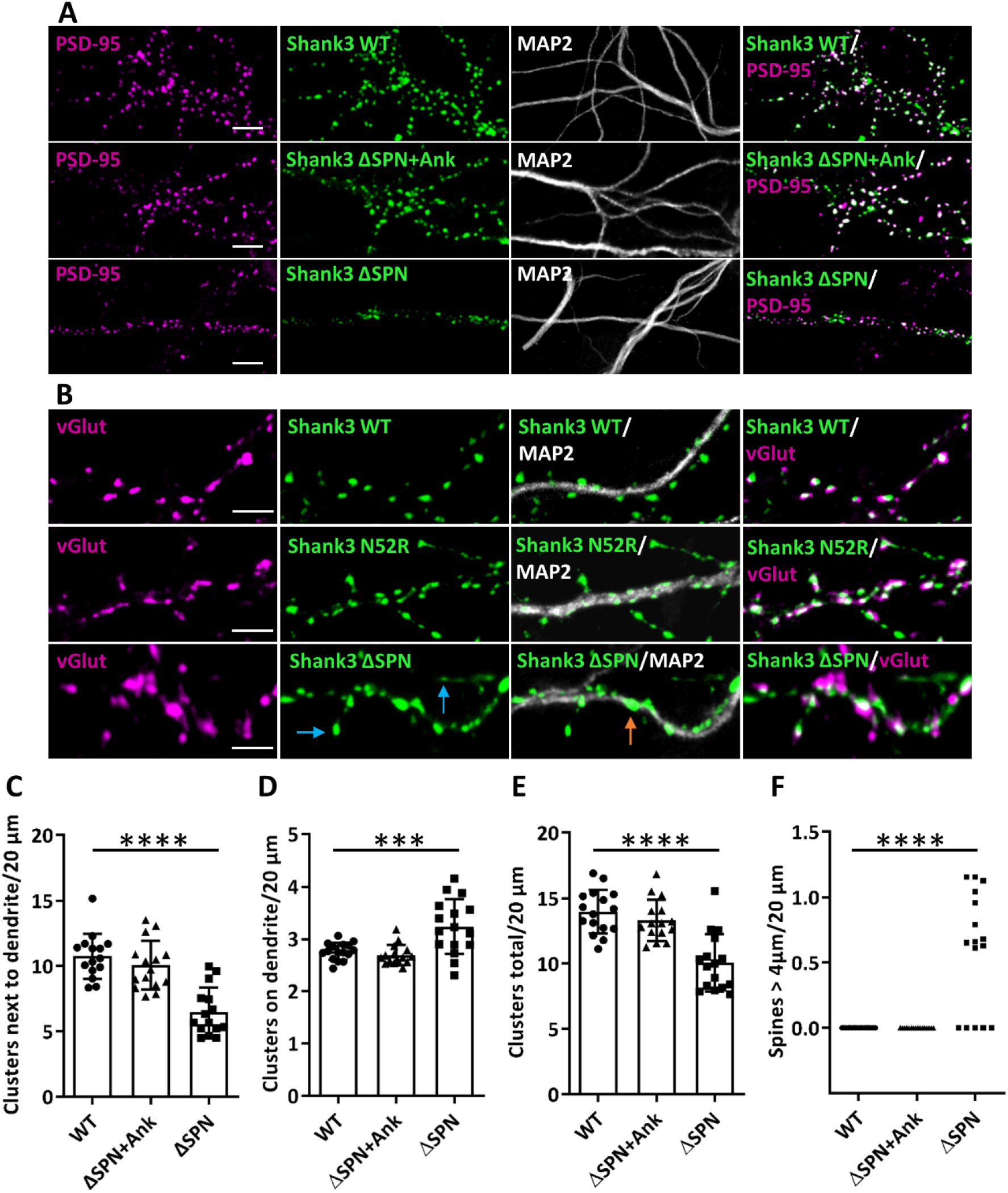
The ΔSPN variant, but not the ΔSPN+Ank variant, induces irregular Shank3 clusters in neurons. **A**,**B**. Neurons were transfected as in Fig. 7. Cells were costained for MAP2 and the postsynaptic marker PSD-95 (A), or the presynaptic marker vGlut (B). Magnifications of dendritic segments are shown (scale bar 5 µm). Blue arrows point to Shank3 clusters on long, thin spines; orange arrows point to atypical, dendritic shaft associated clusters. The N52R mutant Shank3 is included here for comparison (24). **C-F**. Quantitative evaluation of the number of Shank3 clusters (C-E) and number of Shank3-positive Spines > 4 µm (F) per 20 µm dendrite. ***, ****, significantly different from WT; p<0.001, 0.0001, respectively, analysis of 45 dendrites of n=15 neurons from three independent experiments; one-way ANOVA followed by Dunnett’s multiple comparisons test.

We noted that the pattern observed here for the ΔSPN Shank3 protein is very similar to the distribution observed with another Shank3 variant, namely N52R, an artificial mutant that was shown to induce opening of the SPN-Ank tandem (24). Results for expression of this mutant are included here for comparison (Fig. 8B), again showing Shank3 clusters adjacent to the dendritic shaft and some clusters on thin, long protrusions, as observed before (24). On the other hand, we expressed a Q106P mutant construct. This variant, observed in an autistic patient, alters the linker between SPN and Ank domains and selectively interferes with αCaMKII binding (23). Microscopic analysis of transfected neurons showed a very normal distribution, with most Shank3 clusters localized in a spine like pattern similar to WT Shank3 (Fig. 9). We conclude that the mislocalization of full–length Shank3 in atypical synaptic clusters on dendritic shafts, as well as the formation of long, thin spines, is due to the presence of “open”, unprotected Ank repeats (as observed also in the N52R variant), but is not caused by reduced binding of αCaMKII.

**Figure 9.**
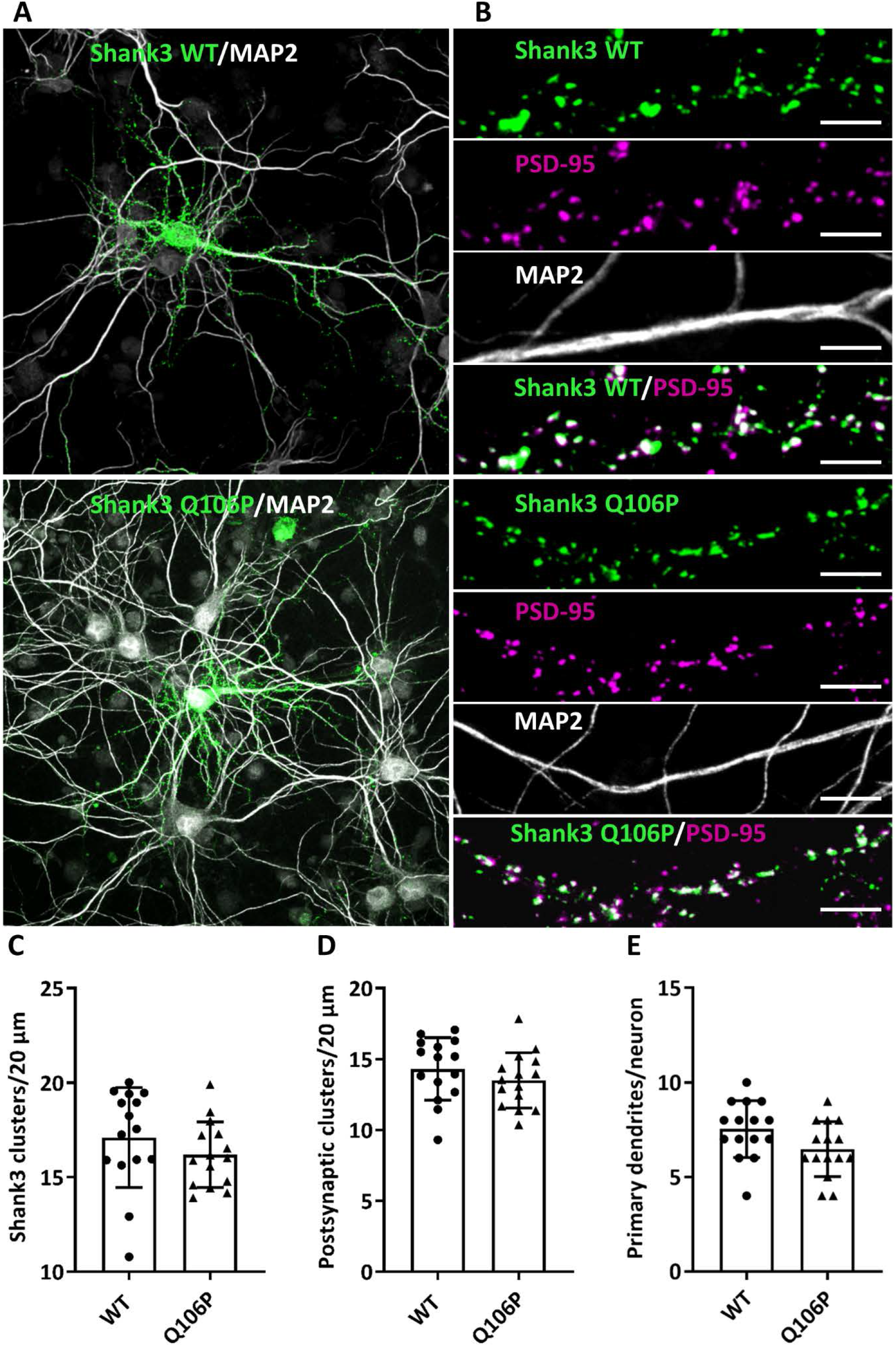
Binding of αCaMKII to Shank3 is not necessary for proper location of Shank3. **A**,**B**. WT and Q106P mutant Shank3 (which is deficient in αCaMKII binding (23)) was expressed in primary hippocampal neurons. Cells were fixed and costained for PSD-95 and MAP2. **A**. Overview images of neurons (scale bar 20 μm). **B**. Magnifications of dendritic segments are shown (scale bar 5 µm). **C+D**. Quantitative evaluation of the total number of Shank3 clusters (C) and the number of postsynaptic Shank3 clusters (D) (Shank3 clusters colocalising with PSD-95) per 20 μm dendrite. ns: non-significant; analysis of 45 dendrites of n=15 neurons from three independent experiments; Student’s t-test. **E**. Quantitative evaluation of the number of primary dendrites per neuron. ns: non-significant; analysis of n=15 neurons from three independent experiments; Student’s t-test.

To further validate this hypothesis, we analysed the stability of the Ank repeat region of Shank3 in the presence and absence of the SPN domain, using the differential scanning fluorimetry (DSF) method. Analysis of the transition curves measured by DSF for the SPN+Ank fragment (1-348), and the Ank only fragment, (aa 99-348), showed that the Ank repeats are less stable than the 1-348 WT fragment. This is represented by a leftward shift in the transition curve and a significant reduction in the mean Tm value of the isolated Ank domain (Figure 10). This indicates that the Ank repeats are stabilised by their association with the SPN domain when in the full 1-348 WT fragment.

**Figure 10.**
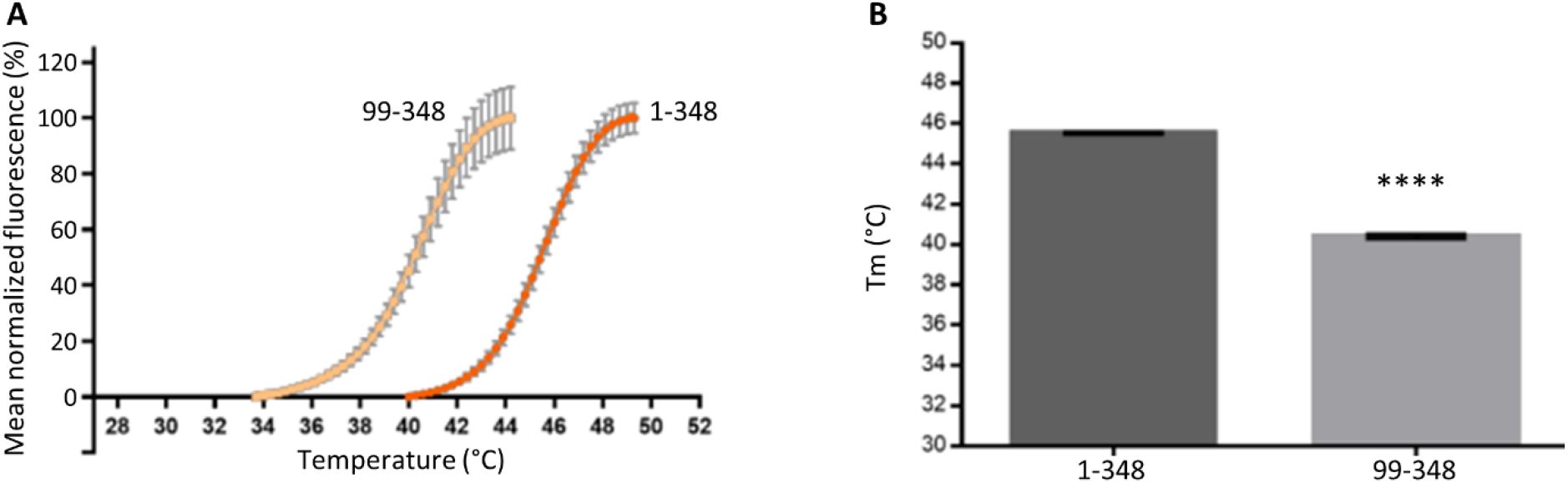
Stability of the Shank3 Ank domain alone is reduced when compared to the SPN+Ank tandem domain. A. Thermal degradation curves measured for the Shank3 1-348 and 99-348 fragments in 0.5 M NaCl buffer, measured as mean normalised fluorescence (%) by DSF (n=3). The thermal transition curve measured for the Ank repeats alone is shifted to the left compared to that of the SPN+Ank curve. B. Calculated Tm values for Shank3 1-348 and ARR 99-348 using transition curves in A. There is a significant decrease in Tm for the ARR 99-348 alone compared to the 1-348 WT fragment (****; unpaired t-test; p<0.0001).

## Discussion

In this study, we attempted to clarify the role of the N-terminal SPN+Ank tandem which is found in all three Shank isoforms at least in one transcript variant. For both Shank1 and Shank3, “long” transcript variants which include these two domains appear to be highly expressed in brain. For Shank2, an SPN+Ank containing transcript variant termed Shank2E is expressed in epithelial cells, whereas Shank2 variants expressed in brain mostly lack both domains. Interestingly, no Shank variants have been reported which lack only the SPN domain or the Ank repeats, suggesting that both domains must occur “in tandem”.

Based on structural analysis of the Shank3 N-terminus, we have initially identified the SPN domain as a binding motif for small G-proteins of the Ras family. Active, GTP-bound forms of both HRas and Rap1a bind the Shank3 SPN domain with high affinity (16, 17). We observed here that coexpression of Shank3 with active HRas or Rap1a in hippocampal neurons can affect the dendritic localization of Shank3; whereas active Rap1a was colocalized with Shank3 in typical spine associated clusters. We noted that active HRas was found in a membrane-associated pattern along dendritic shaft. These data show that small G-proteins may indeed direct Shank3 to specific locations in dendrites; this may be relevant e.g. during synaptogenesis, where small G-proteins can contribute to the formation of postsynaptic clusters. Intriguingly, HRas and Rap1a are associated with Shank3 at different positions on the dendrite. The differential C-terminal modifications are likely to play a role here. Rap1a is geranylgeranylated, whereas HRas is farnesylated; it is known that these modifications target proteins to different cellular microenvironments (34). In addition, it should be noted that Ras and Rap1 activation occur in different, opposing signaling (35).

Active G-proteins bind to a canonical interface on the SPN domain which, in the 3D structure, faces away from the SPN domain (16, 17). In contrast, the αCaMKII binds to the loop connecting SPN and Ank domains and contacts part of the SPN domain. Interestingly, mutation of the conserved Arg12 in the SPN domain of Shank3 (e.g. in the R12C variant found in an autistic patient; (1)), interferes with binding to active G-proteins as well as binding to αCaMKII (16, 22). Coexpression/coprecipitation experiments in 293T cells showed that the N-termini of Shank1 and Shank3, but not Shank2 bound efficiently to αCaMKII. Small G-proteins did not compete with αCaMKII for binding to Shank3. This suggests that the binding sites for Ras/Rap1 and αCaMKII are sufficiently separated from each other to avoid substantial overlap.

We investigated the possibility that Shank proteins, due to their interaction with signalling molecules Ras/Rap and αCaMKII, might alter or regulate specific synaptic signalling pathways. Indeed, WT Shank3 blocked activation of the MAP kinase pathway, whereas the R12C or L68P variants which can not bind active Ras could not do this. However, this did not lead to increased activation of Ras-mediated signalling in brain lysates of Shank3 ko mice.

It was shown before that only the non-phosphorylated, inactive form of αCaMKII can bind to the Shank3 N-terminus (22), whereas another binding site for active, T286-phosphorylated αCaMKII exists in the C-terminal half of the protein (36). Upon preparing PSD fractions from different brain regions of Shank1 as well as Shank3 deficient mice, we observed no change in the absolute amount of PSD-associated αCaMKII. In contrast, the level of phosphorylated αCaMKII is reduced in both mouse lines, indicating that long, SPN+Ank containing Shank variants are negative regulators of αCaMKII activation. Thus, N-termini of brain-expressed long Shank isoforms can alter signalling pathways which have been implicated in NMDA receptor mediated synaptic plasticity.

Finally, we assessed the relevance of the intramolecular interaction within the SPN-Ank tandem domain. Using a full-length Shank3 expression construct, we observed that a GFP-Shank3 fusion protein lacking the SPN domain exhibited an atypical localization in dendrites. Many Shank3 clusters were found on the shaft of dendrites, and some were localized on long, filopodia-like spines extending more than 4 µm from the main dendrite. All of these clusters were positive for PSD-95 and vGlut1, suggesting that these were indeed part of functional synaptic sites. We speculated that this atypical localization of synapses arose due to premature, or uncontrolled aggregation of Shank3 via its “free” Ank repeats. Further support for this concept was obtained by expressing a Shank3 mutant lacking the entire N-terminus (SPN+Ank). Here, localization of the expressed GFP-Shank3 protein became normal again, as clusters were observed in spine like protrusions, whereas shaft-associated clusters were strongly reduced.

Two missense mutants were included here in the context of full-length Shank3 to further clarify the role of the intramolecular SPN-Ank interaction. For the N52R mutant we know that it induces an open conformation of the SPN-Ank tandem. We had previously observed that this leads to irregular clusters on the dendritic shafts (24), and used this mutant here for comparison. As before, we observed irregular, shaft associated clusters. On the other hand, the Q106P variant in the linker between SPN and Ank does not affect the SPN-Ank interaction but selectively reduces interaction with the αCaMKII (23). Full-length Shank3-Q106P was localized in a typical spine associated pattern clearly indicating that interaction with αCaMKII is not necessary for formation of proper Shank3 clusters.

In summary, our data show that the SPN domain, through binding to active Ras family G-proteins, may alter the dendritic localization of Shank3. Simultaneous binding to αCaMKII at the linker between SPN and Ank allows for a negative regulation of the αCaMKII activity at postsynaptic sites. The SPN-Ank tandem must be in a closed conformation not only for binding αCaMKII, but also for preventing Ank-Ank aggregation which may be associated with the mislocalization of synaptic Shank3 clusters on dendritic shafts. Indeed, our measurements of thermal stability of the Shank3 N-terminus show that the SPN domain stabilizes the Ank repeats (Fig. 10). So far, we do not know whether there is any physiological situation where the SPN-Ank tandem appears in an open conformation. A free, or “open” SPN domain binds to F-actin (24), which is likely to be relevant upon maturation of postsynaptic sites. Some kind of trigger for a transition from closed to open conformations is as yet unknown. Active Ras family members might be considered, but they appear to bind efficiently to the closed formation, as seen in Fig. 4 and as observed by (17). Importantly, several mutations found in autistic patients (e.g. R12C; L68P; Q106P; P141A) disrupt the complex interaction pattern at the SPN-Ank tandem, suggesting pathological relevance of open as well as closed conformations.

## Acknowledgements

The authors thank Hans-Hinrich Hönck (UKE Hamburg) for excellent technical assistance. We thank UKE microscopy imaging facility (umif) for providing confocal microscopes. The work was supported by Deutscher Akademischer Austauschdienst (DAAD; to F.H.-N.), Deutsche Forschungsgemeinschaft (DFG; KR 1321/9-1; to H.-J.K) and the Biotechnology and Biological Sciences Research Council (BBSRC) DTP fellowship (to E.W).

